# Live-cell invasive phenotyping uncovers the ALK2/BMP6 iron homeostasis pathway as a therapeutic vulnerability in LKB1-mutant lung cancer

**DOI:** 10.1101/2023.06.14.544941

**Authors:** Junghui Koo, Chang-Soo Seong, Rebecca E. Parker, Bhakti Dwivedi, Robert A. Arthur, Ashok Reddy Dinasarapu, H. Richard Johnston, Henry Claussen, Carol Tucker-Burden, Suresh S. Ramalingam, Haian Fu, Wei Zhou, Adam I. Marcus, Melissa Gilbert-Ross

**Affiliations:** Department of Hematology and Medical Oncology, Emory University School of Medicine, Atlanta, GA, USA; Cancer Biology Graduate Program, Emory University, Atlanta, GA, USA; Biostatistics and Bioinformatics Shared Resource, Winship Cancer Institute of Emory University, Atlanta, GA, USA; Emory Integrated Computational Core, Emory University School of Medicine, Atlanta, GA, USA; Department of Pharmacology and Chemical Biology, Emory University School of Medicine, Atlanta, GA, USA; Department of Neurology, Emory University School of Medicine, Atlanta, GA, USA

## Abstract

The acquisition of invasive properties is a prerequisite for tumor progression and metastasis. Molecular subtypes of KRAS-driven lung cancer exhibit distinct modes of invasion that likely contribute to unique growth properties and therapeutic susceptibilities. Despite this, pre-clinical discovery strategies designed to exploit invasive phenotypes are lacking. To address this, we designed an experimental system to screen for targetable signaling pathways linked to active early invasion phenotypes in the two most prominent molecular subtypes, TP53 and LKB1, of KRAS-driven lung adenocarcinoma (LUAD). By combining live-cell imaging of human bronchial epithelial cells in a 3D invasion matrix with RNA transcriptome profiling, we identified the LKB1-specific upregulation of bone morphogenetic protein 6 (BMP6). Examination of early-stage lung cancer patients confirmed upregulation of BMP6 in LKB1-mutant lung tumors. At the molecular level, we find that the canonical iron regulatory hormone Hepcidin is induced via BMP6 signaling upon LKB1 loss, where intact LKB1 kinase activity is necessary to maintain signaling homeostasis. Furthermore, pre-clinical studies in a novel Kras/Lkb1-mutant syngeneic mouse model show that potent growth suppression was achieved by inhibiting the ALK2/BMP6 signaling axis with single agents that are currently in clinical trials. We show that alterations in the iron homeostasis pathway are accompanied by simultaneous upregulation of ferroptosis protection proteins. Thus, LKB1 is sufficient to regulate both the ‘gas’ and ‘breaks’ to finely tune iron-regulated tumor progression.

## Introduction

Tremendous progress has been made towards treating KRAS-mutant cancers, including the discovery of both direct and indirect targeting strategies(1). Despite these advances, objective response rates for currently approved targeted therapies remain low. KRAS-driven lung cancers are particularly challenging due to the presence of heterogeneous tumor suppressor landscapes that confer unique biology and compensatory pro-growth and survival pathways(2). Recent experimental studies showing variable responses and intrinsic resistance to direct KRASG12C inhibition(3,4) indicate the necessity of incorporating co-occurring tumor suppressor mutations into clinically relevant screening approaches.

Comparative gene expression analysis has been used successfully to discover novel biological aspects of heterogeneous KRAS-mutant lung cancers, including those carrying either *TP53* or *LKB1* mutations(5,6). Integrative approaches using mouse and human tumor cell lines has led to discoveries for novel targeted therapeutic vulnerabilities, particularly for LKB1-mutant patients who are largely resistant to standard-of-care and immunotherapies(7). Efforts to target pro-growth kinase cascades have been met with mixed results(8). One possible explanation for this is the inability of two-dimensional cell culture models to accurately mimic the three-dimensional physiology of lung tissue(9) – and hence fail to predict the dominant signaling molecules driving both cell growth and proliferation during cell:cell and cell:matrix interactions.

In order to identify alternative pro-growth targets that function within a social cell biological context, we took a novel approach that relied upon 3D live-cell phenotyping coupled with comparative gene expression analysis of engineered genetic subsets of invasive human bronchial epithelial cells (HBECs). Using this approach, we identified BMP6 signaling as uniquely upregulated in partially transformed and invasive LKB1-mutant spheroids. To show clinical relevance we probed gene expression data from early-stage lung cancer patients to show that *BMP6* expression is highly elevated in LKB1-mutant lung cancer patients. We coupled this approach with pre-clinical studies in a novel Kras/Lkb1-mutant syngeneic mouse model to show that single-agent potent growth suppression was achieved by inhibiting the ALK2/BMP6 signaling axis with agents that are currently in clinical trials. Thus, image-guided 3D phenotyping is a powerful new tool to advance clinically relevant and personalized targeted treatment strategies that have the potential to achieve single-agent complete responses for KRAS/LKB1-mutant lung cancer patients.

## Results

### Generation and characterization of a panel of subtype-specific human bronchial cells for invasive phenotyping

To generate a series of isogenic cell lines that model clinically relevant early invasion events, immortalized HBEC3-KT cells were stably depleted for LKB1 (L) and TP53 (P) both alone and in a mutant KRAS^G12D^ background (Fig 1A). These isogenic lines include Control (C), KRAS^G12D^ (K), KRAS^G12D^/shRNA-LKB1 (KL), and KRAS^G12D^/shRNA-TP53 (KP) lines, which were validated using western blotting (Fig.1B). To begin to evaluate in vitro transformation phenotypes related to invasion we assessed the expression of epithelial-mesenchymal transition (EMT) markers, E-cadherin and vimentin. E-cadherin was greatly reduced in KL and KP cells, and vimentin was increased. Analysis of cell cycle phasing showed an increase in the percentage of cells in the G2/M phase with the highest percentage in KL and KP cells (Fig. 1C). K, KL and KP lines showed an increase in cell growth rates (Fig 1D). To characterize 2D migration and invasion phenotypes, transwell assays with and without invasion matrix were performed. Similar to the cell growth and cell cycle assays above, KL and KP lines had significantly increased 2D migration and invasion compared to control cells (C, L, and P, and K) (Supplemental Fig. 1 A-D). These data suggest that KL and KP cells are partially transformed in vitro and can be used to screen for targetable pathways linked to early invasion.

**Figure 1.**
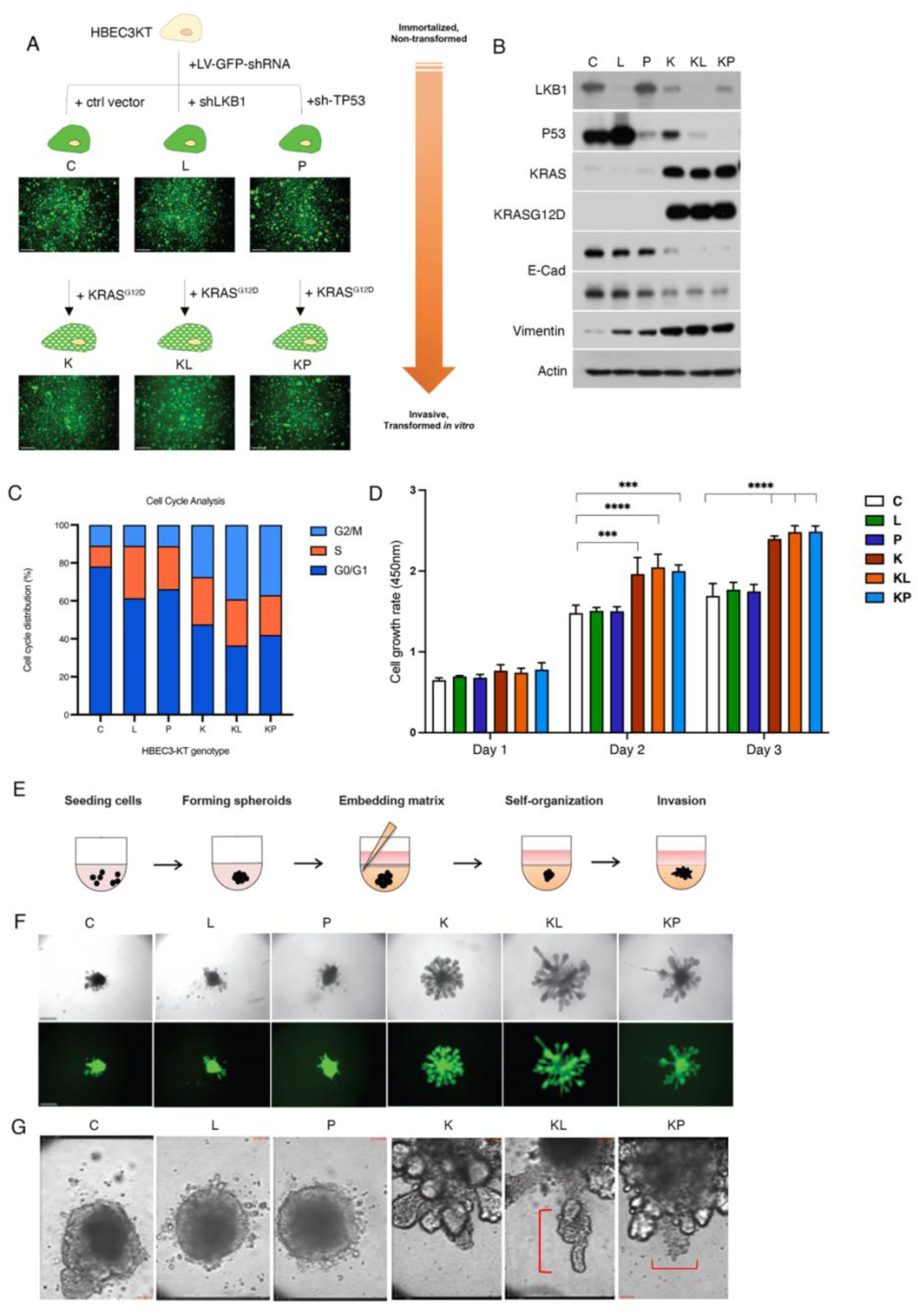
Generation and characterization of in vitro transformation phenotypes and 3D invasive phenotyping. A) Schematic depicting knockdown and overexpression strategy to generate isogenic subsets of HBEC cells for invasive phenotyping. Scale bar = 200µm B) Western blot to verify tumor suppressor, oncogene status and EMT markers in isogenic HBECs. C) Graph depicting quantitative analysis of cell cycle distribution from each isogenic HBEC genotype. D) Graph depicting mean cell proliferation rate from isogenic HBECs over the course of 72hrs. Error bar represents ± SEM of data obtained from quadruplicate samples. E) Schematic representation of 3D invasive phenotyping assay. F) Bright-field and corresponding fluorescent images of live-cell 3D phenotype of isogenic HBECs 4 days post-embedding. Scale bar represents 200 μm. G) Representative still images of focal point time-lapse microscopy (36hr post-embedding) show protruding cell clusters at the leading branching edge, followed by budding formation or ductal elongation (red line). Scale bar = 16 μm.

### KL and KP isogenic HBEC3KT spheroids exhibit distinct autonomous invasion phenotypes

To establish a 3D platform for invasive phenotyping, we created a mixture of Matrigel and Collagen I that supported the 3-D morphogenesis and invasion of partially transformed HBEC3-KTs without the need for feeder fibroblasts (Fig. 1E). Initially, the K, KL and KP cells formed partially aggregated loose spheroids but over the course of several days developed budding and branching morphologies consistent with lung morphogenesis (Fig. 1F). We continued to monitor the behavior of spheroids over several days using live-cell imaging. KP spheroids contained unique lobular structures that continued to grow, spin, and display single invading cells into the surrounding matrix, while KL spheroids exhibited strong collectively invading cellular strands (Fig. 1G, Supplemental Fig. 2, and Supplemental Movie File 1). K spheroids continued to exhibit branching morphogenesis but no invasion, and C, L and P spheroids remained in loose (C) or tightly associated form (L and P). Taken together, these results confirm that our subtype-specific spheroids exhibit unique and cell-autonomous growth and invasion phenotypes in our 3D platform.

### Transcriptomics of subtype-specific invasion reveals upregulation of BMP6 in LKB1-mutant LUAD

To investigate differential RNA expression in spheroids during early invasion, we isolated RNA from each isogenic spheroid 7 days post-embedding in 3D matrix, the point at which K, KL and KP spheroids exhibited either morphogenesis or distinct invasion phenotypes. RNAseq was performed and differentially expressed genes (DEGs) were identified for each comparison of interest with respect to the Control (Ctrl) line. There was a total of 984 significant DEGs (up = 583; down = 401) from the KL versus KP comparison (Fig. 2A). To identify potential upregulated genes that are specific to *LKB1* loss, we analyzed genes that were upregulated in KL vs. K, but not in K vs. C (Fig. 2B), which yielded 207 candidate genes. We further narrowed our candidates to 32 potential targetable hits that were significantly upregulated in the KL vs. KP (in comparison to C) that were uniquely upregulated in KL vs. K (Fig. 2C). From this list we probed the expression of candidate genes from TCGA data in genetic subtypes of human lung adenocarcinoma. Our analysis revealed that BMP6 is upregulated in LKB1 mutant lung adenocarcinoma when compared to other molecular subtypes of LUAD (Fig. 2D). BMP6 is a ligand of the Transforming growth factor β (TGFβ) superfamily and plays an important role in iron metabolism {reviewed in Fisher.2022}. Iron induces treatment-resistant cancer stem cells and aggressive phenotypes in lung cancer cells(10), therefore, we chose to pursue BMP6 and validated upregulation of expression in our KL HBECs (Fig. 2E). Taken together, our data show that 3D invasive phenotyping coupled with transcriptomics can be used to identify clinically relevant targets in LKB1-mutant cancers.

**Figure 2.**
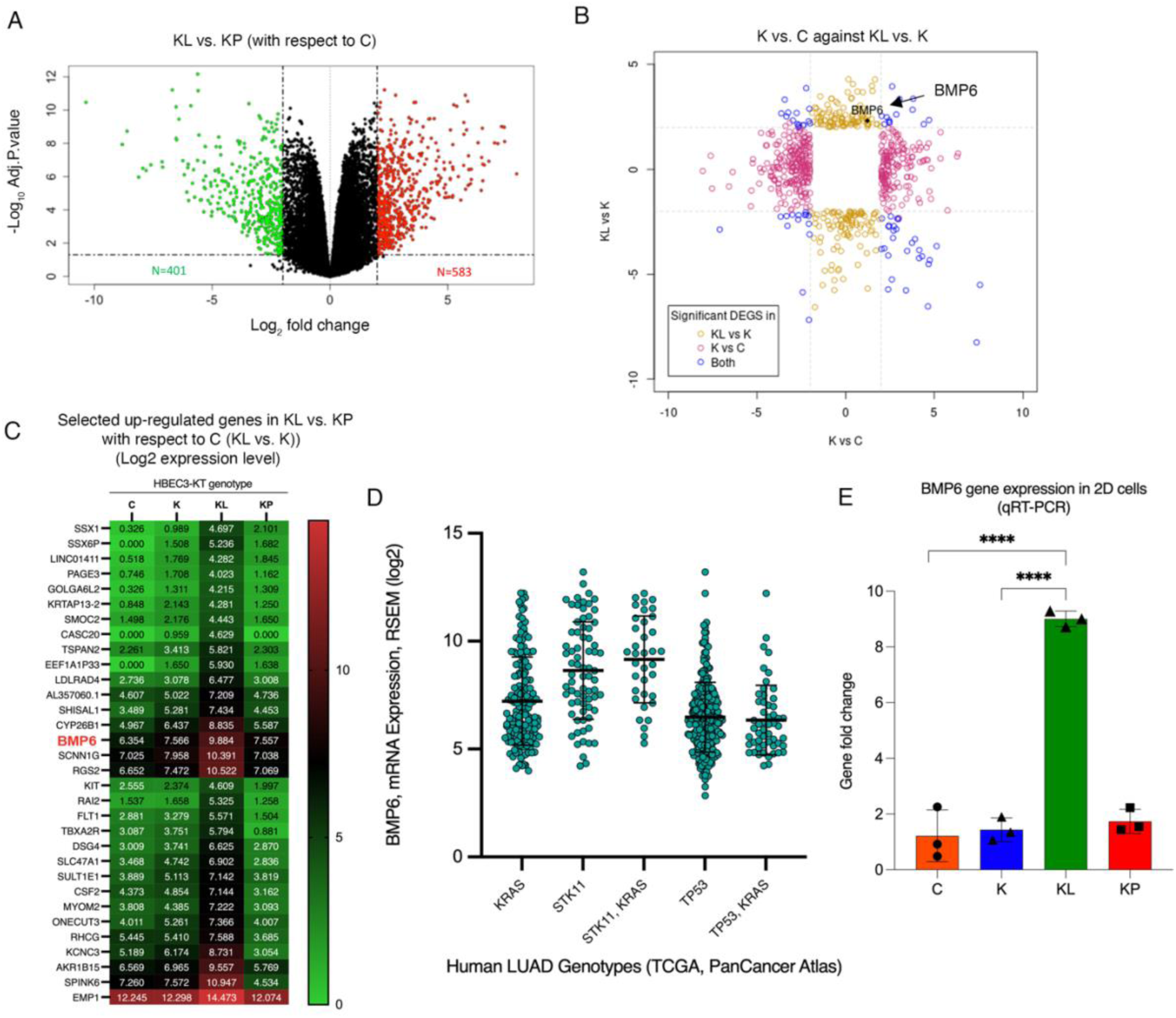
BMP6 expression is uniquely upregulated in response to loss of LKB1 in invasive HBECs and lung adenocarcinoma patients. Volcano plot depicting significantly differentially expressed genes (upregulated, N = 583 and downregulated, N = 401) genes between invasive KRAS/LKB1 (KL) vs. KRAS/TP53 (KP) 3D spheroids with respect to non-invasive control (Control) HBEC spheroids. B) Scatterplot of differentially expressed genes (absolute fold change of >= 2 or more and FDR <= 0.05) showing K vs. C against KL vs. K shown by significance in either or both comparisons. C) Heatmap depicting mean log2 transformed expression levels ofselected differentially upregulated genes (KL>KP and KL>K>C) between isogenic 3D spheroids. D) Graph generated using cBioPortal depicting mean BMP6 mRNA expression from genetic subtypes of lung adenocarcinoma (LUAD) patients. Each circle represents an individual patient sample. Error bars represent standard deviation. E) Graph depicting fold change in BMP6 gene expression among isogenic HBECs.

### LKB1 restricts BMP6 expression and BMP Type I receptor signaling using a kinase-dependent mechanism

BMP6 ligand activates Activin A receptor-type I (ALK2) and is a critical regulator of Hepcidin expression and iron balance in normal and cancerous tissues(11,12). Despite this knowledge, the link between LKB1, iron homeostasis and BMP6 signaling in cancer cells is unexplored. To probe BMP6 function in KRAS/LKB1-mutant lung cancer, we first assayed the expression of BMP6 protein from 3D HBEC spheroids by immunofluorescence and found high levels in KL cells relative to other isogenic subtypes (Fig. 3A). We performed Western analysis from 2D and 3D HBEC cells which showed an increase in intracellular and secreted BMP6 ligand in KL HBECs (Fig. 3B). To test autonomous activation of BMP6 signaling we tested Hepcidin expression, which was greatly increased in KL 3D lysates (Fig. 3B). To test whether LKB1 negatively regulates BMP6 signaling through the canonical Smad1/5/8 signaling pathway we restored wild-type LKB1 wildtype (LKB1-WT) in LKB1-deficient H157 NSCLC cells. BMP6-Smad1/5/8 pathway-mediated gene targets including ID-1, ID-3 and BMP6 itself were decreased by exogenously restored wild-type LKB1 (Fig. 3C). We expressed kinase-dead LKB1 (LKB1-K78I) in H157 cells which was unable to inhibit BMP6-Smad1/5/8 signaling. These data suggest that LKB1 negatively regulates the BMP6-Smad1/5/8-regulated iron homeostasis pathway using a kinase-dependent mechanism.

**Figure 3.**
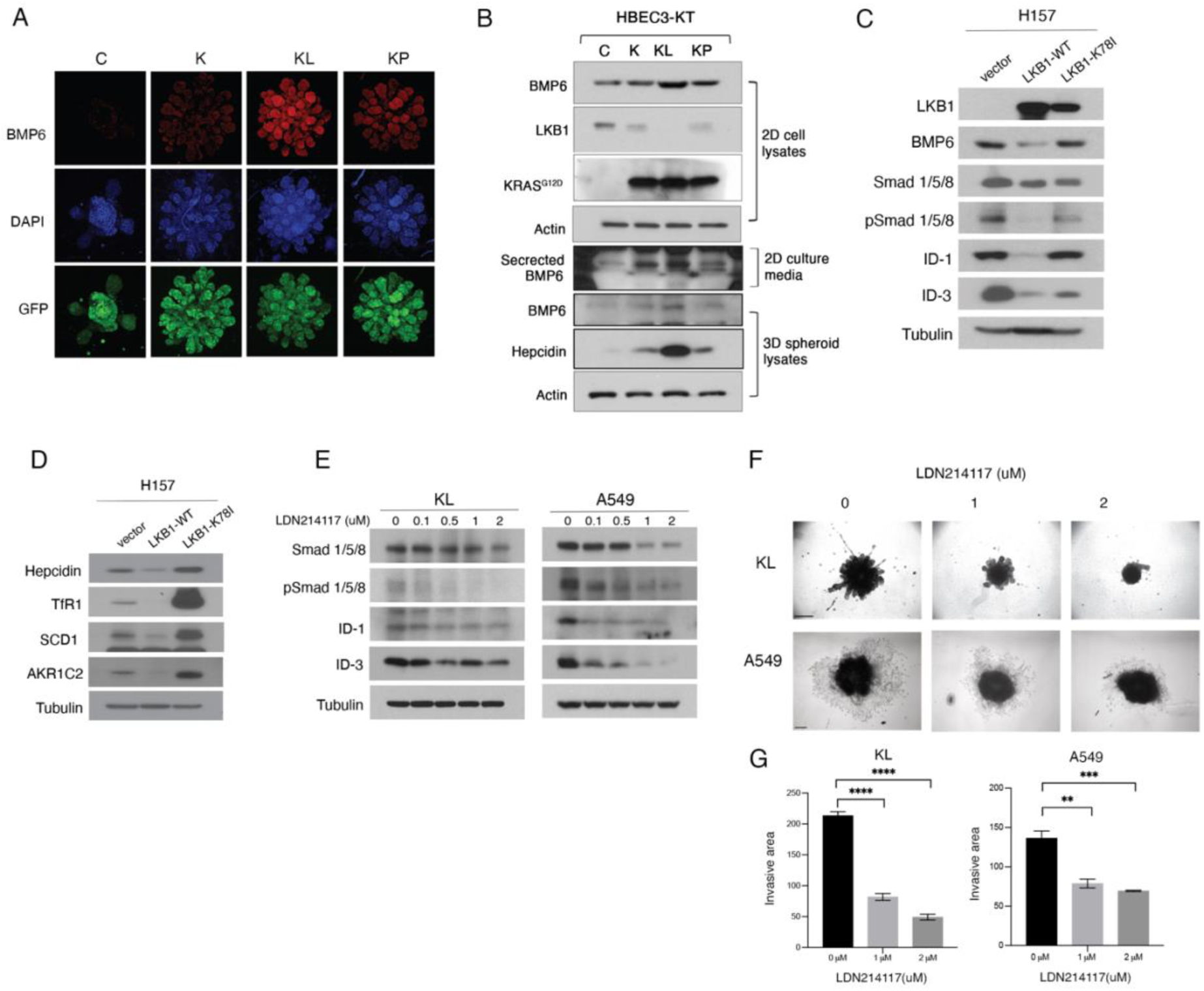
LKB1 negatively regulates the BMP6-induced iron homeostasis pathway through hepcidin and pharmacological inhibition of ALK2 inhibits invasion of KL cells in vitro. A) Confocal images of immunofluorescence for BMP6 protein expression (red) in 3D HBECs 3D of the indicated genotypes (DAPI labels nuclei and all cells express cytoplasmic GFP). Western blot of the indicated proteins detected from 2D and 3D lysate and media from HBECs of the indicated genotypes. C) Western blot of BMP6-regulated Smad signaling components in LKB1-null H157 cells that express vector control, wild-type LKB1 (LKB1-WT) or kinase dead LKB1 (LKB1-K78I). D) Western blot of iron homeostasis and ferroptosis protective genes in H157 cells expressing vector control, LKB1-WT or LKB1-K78I. E) Western blot analysis of BMP6/ALK2-regulated Smad signaling in KL HBECs and A549 (LKB1-null) cells treated with increasing concentrations of LDN214117 for 24hrs. F) Brightfield images of 3D spheroids of the indicated genotypes treated with the indicated concentration of LDN214117 and embedded in invasion matrix for 72hrs. G) Quantitation of invasive area in for LDN214117-treated spheroids. (graph depicts mean of 3 biological replicates).

In cancerous tissues, the expression of excess Hepcidin results in accumulation of intracellular iron (Fe2+) pools to promote cancer cell growth and proliferation(12). Hepcidin accomplishes this by directly promoting the degradation of ferroportin receptor (FPN), which acts to export excess intracellular Fe2+. This is accompanied by an increase in the expression of the transferrin receptor (TfR1), which acts to import ferric iron (TF-Fe3+) into the cell. To test whether the increased Hepcidin in LKB1-mutant cells is functional, we expressed LKB1-WT in H157 cells which resulted in the downregulation of both hepcidin and the TfR1. The ability of LKB1 to downregulate hepcidin and TfR1 was dependent on its kinase activity (Fig. 3D).

Cancer cells balance their metabolic need for iron with protection from iron overload and cell death. Recent studies have linked LKB1 and KEAP1 co-mutations with the regulation of ferroptosis, an iron-dependent form of cell death(13). To test whether ferroptosis protective genes are co-regulated by LKB1 alongside the BMP6-hepcidin iron homeostasis pathway, we expressed LKB1-WT and LKB1-K78I in H157 cells and tested the expression of stearoyl-CoA desaturase 1 (SCD1) and Aldo-Keto Reductase-1C (AKR1C2) proteins. Expression of LKB1 in H157 cells inhibited both SCD1 and AKR1C2 in a kinase-dependent manner (Fig. 3D). These data suggest that in the presence of wild-type KEAP1, LKB1 can simultaneously regulate both iron homeostasis and ferroptosis protection pathways.

### Inhibiting BMP6/ALK2 receptor signaling decreases 3D invasion in KL cells and reduces tumor growth in an immunocompetent mouse model

ALK2 is an oncogenic driver in 25% of pediatric diffuse intrinsic pontine glioma (DIPG) tumors, but its role in lung cancer is largely unexplored. Several small molecules target BMP type I receptors including LDN214117, which is a small molecule kinase inhibitor that exhibits a high degree of selectivity for BMP6-activated ALK2(14). Increasing concentrations of LDN214117 resulted in a dose-dependent inhibition of SMAD 1/5/8 activation and downstream target gene induction in KL HBECs and LKB1-deficient A549 lung cancer cells (Fig. 3E). The same cell lines were then utilized in 3D invasion studies with LDN214117 treatment, which significantly inhibited invasion in both cell types (Fig. 3F, G).

To test whether the pharmacological inhibition of ALK2 could be used as an in vivo treatment strategy, we created two mouse tumor cell lines from the Kras^G12D^/Lkb1^fl/fl^ genetically engineered mouse model (KL GEMM). JK43-P and JK43-M were generated from a primary lung tumor and a tumor-positive mediastinal draining lymph node, respectively (Fig. 4A). The status of oncogene and tumor suppressor expression, and epithelial identity was confirmed by western blot (Fig. 4B). We tested pre-treatment levels of BMP6/ALK2 pathway activation and found that target gene expression was higher in the JK43-M cells compared to the JK43-P cells. However, treatment of both cell lines with the ALK2 inhibitor LDN214117 resulted in dosage-sensitive pathway inhibition (Fig. 4C). In order to test whether treatment with LDN214117 could suppress iron homeostasis and ferroptosis protective pathways in mouse tumor cells we treated JK43-M cells with 1µM of LDN214117 for up to 72h. Iron homeostasis (Hepcidin and TfR1) and ferroptosis-protective proteins (SCD1 and AKR1C2) were decreased by pharmacological inhibition of ALK2 (Fig. 4D). Taken together, these data show that inhibition of BMP6/ALK2 activity in KRAS/LKB1-mutant lung cells is sufficient to inhibit 3D invasion in both early-stage partially transformed cells and an aggressive lung cancer cell line. In addition, our results demonstrate that the JK43-M mouse tumor cell line can be used as a pre-clinical platform for testing therapeutic efficacy in vivo.

**Figure 4.**
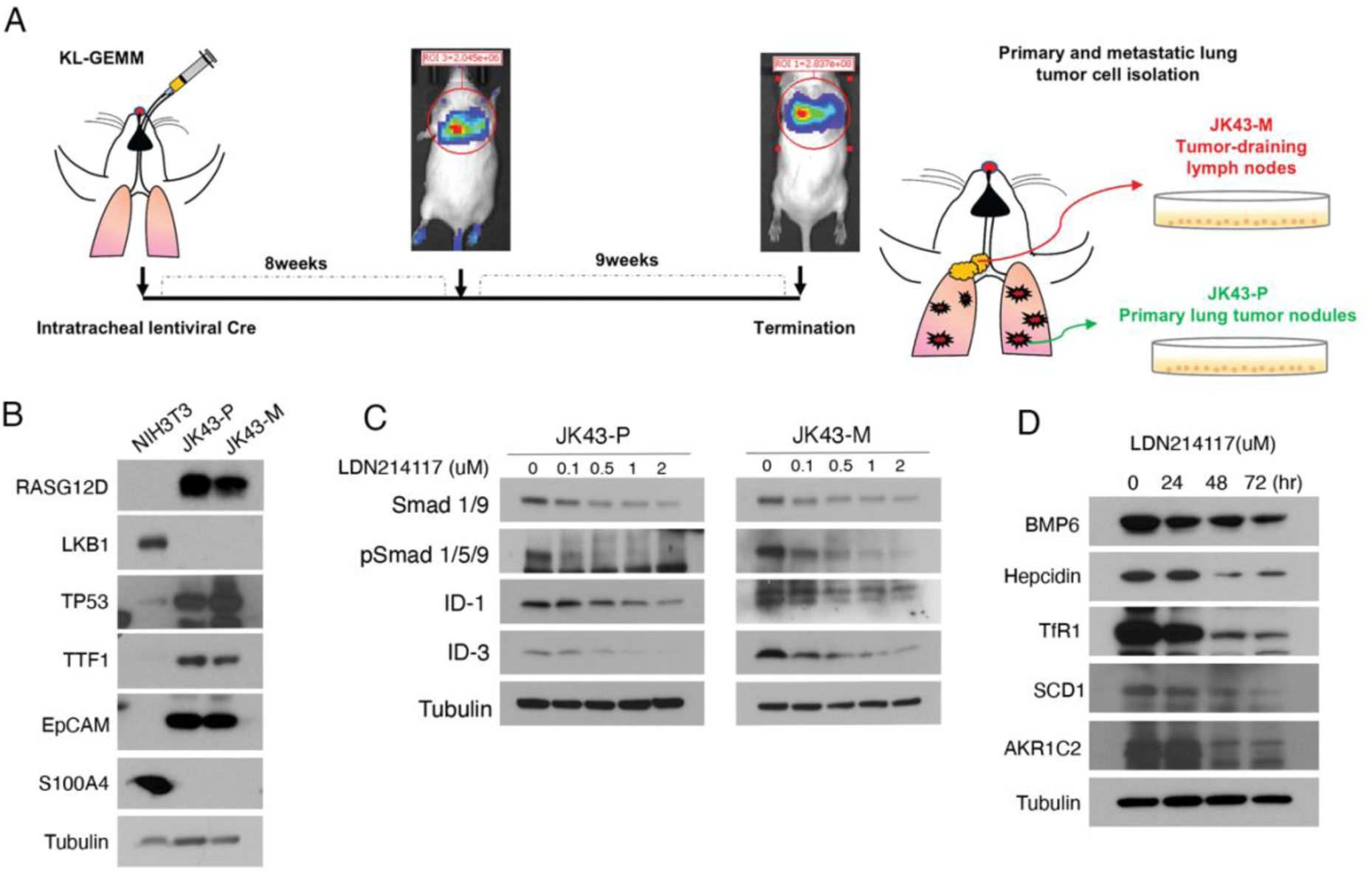
Kras^G12D^/Lkb1^fl/fl^ mouse tumor cells lines respond to AKL2 inhibition by downregulating BMP6-regulated iron homeostasis and ferroptosis protective proteins. A) Schematic of mouse KL tumor cell line isolation. B) Western blot to assess indicated proteins in NIH3T3 (mouse fibroblast cell line), JK43-P and JK43-M tumor cells. C) Western blot of indicated Smad signaling proteins from JK43-P and JK43-M cells treated with increasing concentration of LDN214117. D) Western blot of indicated iron homeostasis and ferroprotective proteins from JK43-M cells treated with 1uM of LDN214117 for 24, 48 and 72hrs.

To determine the *in vivo* efficacy of targeting ALK2 activity with LDN214117 in Kras^G12D^/Lkb^fl/fl^ lung adenocarcinoma, we created a syngeneic mouse model using the JK43-M tumor cell line. JK43-M cells were subcutaneously injected into FVB/NJ mice and treated with 25mg/kg of LDN214117 (Fig. 5A). Tumor volume and weight was significantly inhibited in LDN214117 treated mice compared to the vehicle control (Fig. 5B-D). To explore the functional consequences of direct BMP6 inhibition, we treated KL HBECs and JK43-P and JK43-M cells with a neutralizing anti-BMP6 antibody which resulted in decreased of Smad 1/5/8 signaling and 3D invasion compared with IgG control antibody (Supplemental Figure 3A,B). Lastly, LDN214117 treated tumors exhibited downregulated BMP6 expression and Hepcidin induction (Fig. 5E,F). These data are consistent with a model where LKB1 is sufficient to simultaneously restrict tumor progression via BMP6/ALK2-induced Hepcidin regulated iron homeostasis in addition to ferroptosis protection pathways.

**Figure 5.**
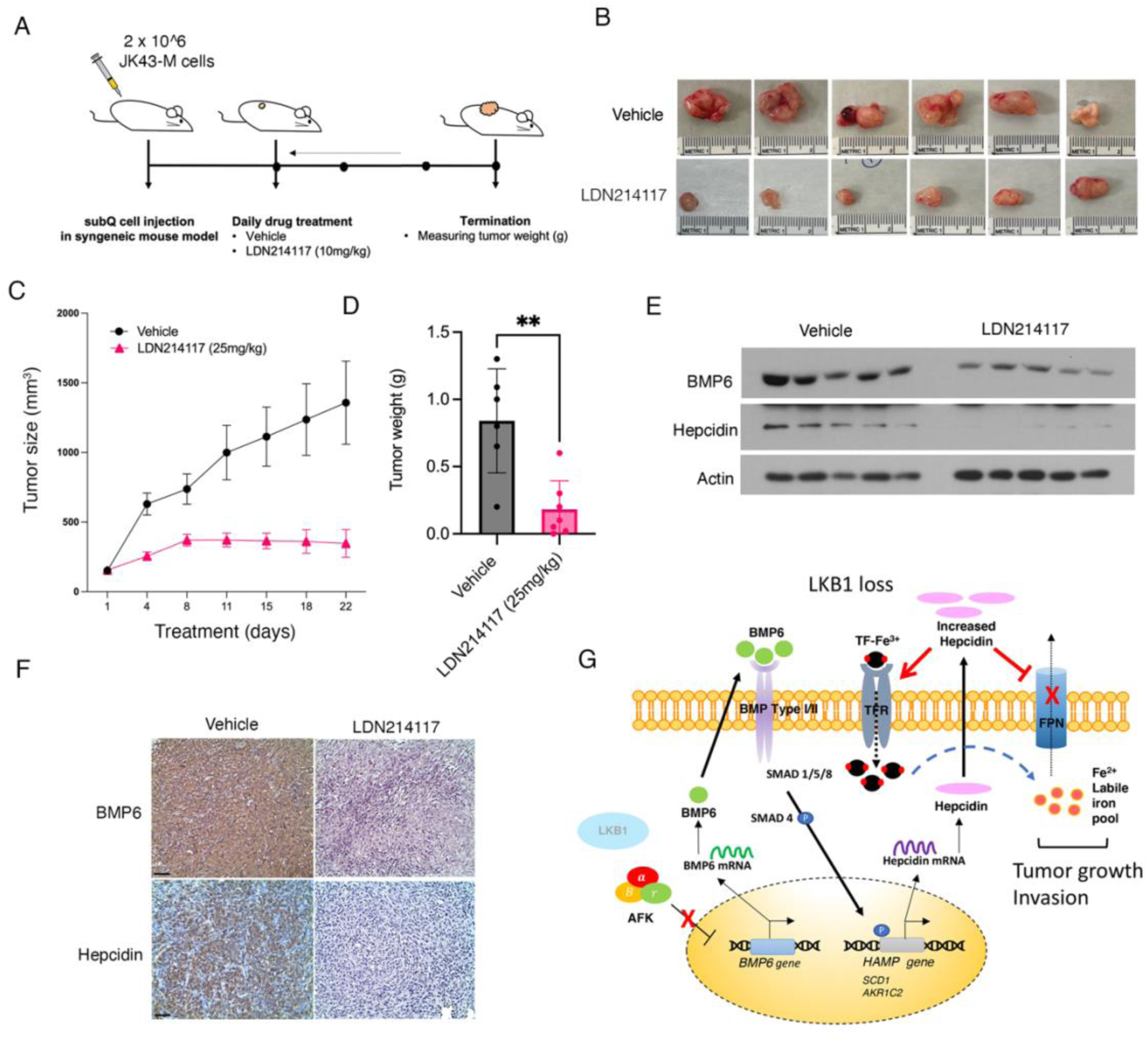
In vivo efficacy of LDN214117 targeting Alk2 in syngeneic mouse model. A) Schematic depicting pre-clinical trial workflow. B) Brightfield images of tumors isolated from vehicle and LDN214117 treated mice. C) Mean tumor volume from mice treated with vehicle (6 mice/group) and LDN214117 (7 mice/group). D) Graph depicting mean tumor weight from vehicle and LDN214117 treated mice. E) Western blot to assess BMP6 and hepcidin expression level in tumors from vehicle and LDN214117-treated mice. F) Representative images of immunohistochemistry for BMP6 and Hepcidin on tumor sections from mice treated with vehicle or LDN214117. F) Model depicting mechanism of altered iron homeostasis in LKB1-mutant tumor cells.

## Discussion

In this study we used live-cell phenotyping as a guide to functional gene expression profiling in a panel of partially transformed HBECs to identify clinically relevant targeting strategies for treatment-resistant LKB1-mutant lung adenocarcinomas. We show that LKB1 loss upregulates BMP6 levels in invasive HBECs and patient tumors, and that LKB1 simultaneously restricts the BMP6/Hepcidin iron regulatory and ferroptosis protection pathways through its kinase activity. Furthermore, we created a novel Kras/Lkb1-mutant syngeneic mouse model using a highly metastatic KL tumor with intact p53. Using this model, we show that tumor invasion and growth can be potently inhibited using a BMP6 monoclonal antibody and the TGFβ-family ALK2 kinase inhibitor. Our results suggest partially transformed epithelial cells grown in 3-D can be used successfully to screen for therapeutic strategies that have the potential to be rapidly translated into the clinic.

We show here that LKB1 is required and sufficient to restrict the BMP6-regulated Smad signaling in a kinase-dependent manner. This is consistent with a previous report showing that LKB1 can negatively regulate the BMP type 1 receptors, including ALK2, in multiple cell lines and in a Drosophila model, including downstream SMAD signaling(15). This report shows a potential protein-protein interaction between LKB1 and ALK2, which could suggest that the LKB1-BMP6 axis characterized here is due to the LKB1 interaction with ALK2 directly, and would further explain why KL cells are sensitive to ALK2 inhibitors in vivo and in vitro. Therefore, when synthesizing the findings here with others, one proposed model is where LKB1 loss potentiates a ALK2-BMP6 receptor ligand interaction that drives downstream BMP6 signaling.

ALK2 is a clinically actionable target in several diseases, and small molecules antagonists are orally bioavailable and well-tolerated in pre-clinical studies with patient-derived models(16). In diffuse intrinsic pontine glioma (DIPG), treatment with ALK2 inhibitors LDN-193189 or LDN-214117 extended survival compared with vehicle control(17). Trials have completed or are ongoing using ALK2 small molecule or biologic inhibitors in different clinical settings(18). Others ALK2 inhibitors, such as Saracatinib, has improved selectivity for ALK2 over ALKs 3, 4, 5, supporting the concept of highly selective ALK2 targeting in the clinical setting(19). Based upon our findings we would propose that the KL genetic sub-type of lung cancer would be uniquely sensitive to this drug class. In addition, the ability of ALK2 inhibitors to penetrate the blood brain barrier suggest they may be efficacious in treating primary lung cancers that have metastasized to the brain. Additional pre-clinical studies to further test this are warranted.

Our data is consistent with a model where LKB1 can apply the ‘gas’ and the ‘breaks’ to finely tune a cancer cell’s need for iron-regulated metabolic growth with protection from iron overload and cell death. Hepcidin regulates iron homeostasis serving as the key molecule overseeing systemic iron absorption, where its dysregulation results in various iron disorders (https://www.ncbi.nlm.nih.gov/pmc/articles/PMC2855274/). The data here show that loss of LKB1 leads to increased hepcidin levels, and is sufficient to restrict BMP6-regulated Hepcidin in a kinase dependent manner. Moreover, we show that LKB1is sufficient to simultaneously restrict ferroptosis protection proteins in the presence of wild-type KEAP1. Our data also show that BMP6 protein levels are sensitive to LKB1 genetic perturbation as well as pharmacological inhibition with the ALK2 inhibitor. This suggests that changes in BMP6 protein levels may be exploited in the future in KL patients as a positive predictive biomarker of response to ALK2 inhibition.

## Materials and Methods

### Reagents and antibodies

LKB1 (#3050 for human, #3047 for mouse), p53 (Rodent specific, #32532), RASG12D (#14429), E-Cadherin (#3195), Vimentin (#3932), p-Smad1/5/9 (#13820), ID-3 (#9837) and AKR1C2 (#13035) antibodies were purchased from Cell Signaling Technology. BMP6 (ab155963), Smad 1/9 (ab108965), ID-1 (ab192303), Transferrin receptor (ab84036), SCD1 (ab236868), Ferritin (ab75973), EpCAM (ab71916) and TTF1 (ab76013) antibodies were from Abcam. KRAS (sc-30), p53 (sc-126 for human), a-Tubulin (sc-8035) antibodies were from Santa Cruz Biotechnology. Hepcidin Antimicrobial Peptide antibody (NBP1-59337) was from Novus Biologicals. Actin (A2066) was from Sigma-Aldrich. Horseradish peroxidase (HRP)-conjugated secondary antibodies (Jackson ImmunoResearch) were used for western blotting.

Neutralizing BMP6 antibodies (MAB507 for human, MAB6325 for mouse) were purchased from R&D system. LDN214117 (S7627) for in vitro and in vivo assay was from Selleckchem, and Cell Counting Kit-8 (CK04-11) was from Dojindo.

### Cells culture conditions and viral transductions

The HBEC3-KT cell line was kindly provided from Dr. John Minna (The University of Texas Southwestern Medical Center, Dallas, TX). HBEC3-KT cells were cultured with Keratinocyte serum-free medium (K-SFM; Life Technologies, Gainthersburg, MD) containing 50ug/ml of bovine pituitary extract (BPE; Life Technologies) and 5ng/ml EGF (Life Technologies) as described previously. Custom-designed lentiviral shRNA-LKB1(target seq; 5’-GACAACATCTACAAGTTGT-3’) and shRNA-p53 (target seq; 5’-ACATTCTCCACTTCTTGTT-3’) vector expressing GFP plasmids were purchased from ATCGbio Life Technology Inc, British Columbia, Canada). pLenti6-KRAS^G12D^ lentiviral vector was a gift from Dr. John Minna. Large-scale production of high-titer lentivirus was generated using Lenti-X™ 293T Cells (TaKaRa, #632180) according to an established. Transduced cells were selected with hygromycin B (20ug/ml) and blasticidin (2ug/ml). Stable LKB1 or p53 knockdown and moderate KRAS^G12D^ expression by lentivirus was introduced stepwise manner as described in Fig.1 A and confirmed by western blot.

H157 cells stably expressing LKB1 (LKB1-WT) and kinase-dead LKB1 domain (LKB1-K78I) were generated as described previously(20). Stably knockdown LKB1 H1299 cells were generated by pLKO.1-shRNA LKB1. Transfected cells were selected with 2 ug/ml puromycin. LKB1 deletion was verified by western blot.

BMP6 shRNA (TRCN0000432077) was purchased from Sigma-Aldrich. Stable knockdown BMP6 JK43-P and JK43-M cells were selected with 5ug/ml puromycin.

H157, A549 and H1299 human NSCLC cells, JK43-P and JK43-M KL-GEMM cells were cultured in Roswell Park Memorial Institute (RPMI-1640) media or Dulbecco’s Modified Eagle Medium (DMEM) with 10% fetal bovine serum and 100 units/mL of penicillin and streptomycin and maintained at 37 °C and 5% CO2.

### Western blot analysis

3D cell lysates were extracted from the 3D spheroids after 3D invasion assay in ultra-low attachment 96well plate. Culture media on the top of the matrix was removed and washed in cold PBS and the matrix was depolymerized with 200ul of Cultrex organoid harvesting solution (R&D systems) at 4°C for 2 h and then the isolated cell spheroids were lysed with 1% NP-40 cell lysis buffer containing protease and phosphatase inhibitor cocktail (Thermo-Scientific, #78442) and by microtip sonicator.

For detecting BMP6 and hepcidin from mouse tumor tissues, isolated subcutaneous tumors were minced and lysed in RIPA buffer containing protease and phosphatase inhibitor cocktail by mechanical homogenizer. Samples were centrifuged at 12000rpm for 10min at 4°C and then the supernatant (lysate) was transferred to new tube.

The Bradford protein assay (BioRad Laboratories Inc. CA, USA) was used to measure the concentration of total protein in samples.

#### Cell cycle analysis

Cells were harvested and fixed in 70% cold-ethanol at −20°C for overnight. Cells were washed in PBS and stained with DAPI (4 μg/mL DAPI, 0.25% Triton-X 100 in 1× PBS). DNA content was measured by flow cytometry with FACSCalibur (BD Biosciences), and data were analyzed to determine the distribution of cells with sub-G1, G1/G0, S, and G2/M peaks using FlowJo software.

### Cell growth and viability assays

For examining cell proliferation rate of the transformed HBEC3-KTs, the cells were seeded at the density of 4000 cells/well in 96-well plates in four replicates and cell growth rate was determined using CCK8 (Dojindo) at the indicated time point according to the manufacturer’s instructions. For checking cell viability after treating small molecule, LDN214117, 3000 of cells were seeded in 96-well culture plates in four replicates and treated the next day with the given concentration of LDN214117. Viable cell numbers were determined using a sulforhodamine B (SRB) assay.

## 2D cell migration and invasion assays

For 2D cell migration assay, 1 x 10^4^ cells were seeded in growth factor free-KSFM media in the top transwell chamber (#353097, Corning Inc., NY, USA) and allowed to migrate along a concentration gradient through a polycarbonate membrane with 8 µm pores to the bottom chamber containing KSFM medium with 100ug/ml of BPE, 10ng/ml of EGF and 10% of FBS. After 24h, cells were fixed in 4% formalin and stained with Crystal violet. The stained cells were counted from the 4 randomly chosen fields on the bottom surface of the membrane under 20x magnification of IX-51 microscope. For 2D cell invasion assay, the transwell membranes were coated with Growth Factor Reduced Matrigel (#356231, Corning Inc., NY, USA) for 30min at 37°C incubator and 1.5 x 10^5^ cells were seeded in growth factor free-KSFM medium in the top chamber. Cells were allowed to migrate for 48h across Matrigel-coated membranes to the bottom chamber in KSFM medium containing 100ug/ml of BPE, 10ng/ml of EGF and 10% of FBS. Cells were fixed, stained with Crystal violet. Each sample was triplicated.

## 3D spheroid invasion assays and live cell imaging

For the transformed HBEC3-KTs spheroid invasion assay, 6000 cells in 100ul of K-SFM medium were seeded in 96-well clear round bottom ultra-low attachment microplate (#7007, Corning Inc.) and centrifuged at 200 x g for 3 min in a swinging bucket rotor. When cells assemble loose-aggregated spheroid at 37°C incubator in 4∼5 days after seeding, 50ul of the medium was removed carefully. Matrigel (#356237, Corning Inc.) and rat tail Type I Collagen (#354249, Corning Inc.) mixed invasion matrix was embedded directly into well on ice and then centrifuged at 300 x g for 5 min in a swinging bucket rotor. 100ul of K-SFM medium was added on the top of the matrix after gel polymerization at 37C incubator for 1h and then allowed to invade. Images were taken using Olympus IX51 microscope 10× (0.30 NA air) with an Infinity2 CCD camera. Live cell imaging was performed a Perkin Elmer Ultraview.

For human NSCLC cells and KL-GEMM cells, spheroids were formed in ultra-low attachment 96-well round bottom plates and embedded in invasion matrix in a 35 mm glass bottom dish (Cellvis) and allowed to invade for up to 72 h incubated at 37°C. Images were taken using an Olympus IX51 microscope 10×.

For neutralizing antibody treatment, antibodies were pre-treated in the medium 48h before embedding invasion matrix and added in the media on top of the matrix. For small molecule treatment, LDN214117 was added directly to the invasion matrix during the embedding process, as well as to the growth media added on top of the matrix.

Invasive area was quantified by measuring the total spheroid area around the outer perimeter in ImageJ/Fiji.

## 3D spheroid immunofluorescence

Preparation of spheroids and immunofluorescence staining were performed as described previously(21). The spheroids were stained with rabbit anti-pYFAK397 (ab39967, Abcam), rabbit anti-Vimentin (3932S, Cell Signaling Technology) and DAPI (D1306, Invitrogen). Secondary antibodies were purchased from ThermoFisher Scientific (Anti-Rabbit Alexa Fluor 647, Anti-Rabbit Alexa Fluor 555 Conjugate). After primary and secondary antibody staining, spheroids were imaged with the Leica TCS SP8 inverted confocal microscope (10×) using 1.0 mm z-stack intervals, line scanning (405 nm DMOD Flexible, 488 nm argon, 561 nm DPSS, 633 nm Helium-Neon), and PMT detectors.

### RNA sequencing from 3D spheroid and gene expression analysis

RNA-seq was performed in triplicate on HBECK3-KT C, L, P, K, KL and KP. Total RNA was extracted from the spheroids at day 7 of 3D spheroid invasion assay using a RNeasy Kit (Qiagen, Valencia, CA, USA). The quantitation, integrity and purity of the extracted total RNA samples were assessed by the Emory Integrated Genomics Core (EIGC) using 2100 Bioanalyzer (Agilent) and RNA sequencing was performed by Novogene, Co., Ltd. Data processing, quality control, read alignment and statistical analyses were performed by the Emory Biostatistics and Bioinformatics Shared Resource, as previously described(21). Raw read data (fastq) QC was performed using FastQC v0.11.7. Post-filtered reads were mapped against Ensemble Human GRCh38.p12 release 95 reference genome using the STARaligner v2.7.0e. Expression quantification was obtained using *featureCounts*. Normalization and pairwise differential analysis were determined using median-ratios method in DESeq2 and log2 transformed. Differentially expressed genes (DEGs) were identified using a moderated t-test as implemented in the Limma R package. Heatmaps were created by unsupervised clustering of log2 transformed normalized expression data for the significant DEGs, and the resulting data were analyzed using a Gene Ontology (GO) enrichment analysis and Kyoto Encyclopedia of Genes and Genomes (KEGG).

### RNA extraction and RT-PCR

Total cellular RNA was extracted from cell monolayers with a RNeasy Kit following manufacturer’s instructions, and then cDNA was synthesized by iScript select cDNA synthesis kit (Bio-Rad; Hercules, CA). Real-time RT-PCR was performed with the iTaq Universal SYBR Green Supermix (Bio-Rad) on 7500 Fast Real-time PCR System (Applied Biosystems) following the manufacturer’s instruction. Primer sequences are as follows:

### BMP6 (human)

forward 5′-CGTGAAGGCAATGCTCACCT-3′,

reverse 5′- CCTGTGGCGTGGTATGCTGT-3′,

### BMP6 (murine)

forward 5′- AAGACCCGGTGGTGGCTCTA-3′,

reverse 5′-CTGTGTGAGCTGCCCTTGCT-3′,

hepcidin (human)

forward 5′-GACGGGACAACTTGCAGAGC-3′,

reverse 5′-GCCTCTGGAACATGGGCA-3′,

hepcidin (murine)

forward 5′-AACAGATACCACACTGGGAA-3′,

reverse 5′-CCTATCTCCATCAACAGATG-3′,

β-Actin (human)

forward 5′-ACGGTGAAGGTGACAGCAGTCG-3′,

reverse 5′-AATGTGCAATCAAAGTCCTCGGC-3′,

β-Actin (murine)

forward 5′-CCGTGAAAAGATGACCCAGA-3′,

reverse 5′-AGGCATACAGGGACAGCACA-3′

### Generation of primary and metastatic lung tumor cells from KL-GEMM

KRAS^G12D^ and LKB1 ^fl/fl^ -genetically engineered mouse model (KL*luc*-GEMM) was generated as previously described(22). Lung tumors were induced by intratracheal intubation of LV-CMV-Cre-GFP (SL100277, SignaGen Laboratories). From five weeks after lentiviral-Cre infection, tumor development was monitored and imaged on an IVIS Spectrum (Perkin Elmer) once a week until the onset of clinical symptoms (hunched posture, tachypnea and weight loss more than 20%), and the mice euthanized according to IACUC guidelines.

Lung tumor nodules and metastatic mediastinal lymph nodes were isolated from the late stage of male KL-mouse (JK-43) after perfusing via the right atrium with 10ml of ice-cold PBS with the use of a 10ml syringe fitted with a 21G needle. The isolated tumor tissues were minced into small pieces and further incubated in Collagenase/Dispase (#07449, STEMCELL Technologies) and elastase (#07453, STEMCELL Technologies) solution at 37°C, 5% CO2 incubator for 16h. 1mg/ml of DNase I (#11284932001, Sigma-Aldrich) was added to cell suspensions for 15min at room temperature (15 ∼25°C), and then cells were was filtered through sterile cell strainers with pore sizes of 70 µm and 40 µm. The final filtrate was centrifuged at 300× g for 3 min at 4°C, and the resulting pellet was suspended in 10 mL of complete culture medium (Dulbecco’s Modified Eagle Medium (DMEM) with 10% fetal bovine serum and 100 units/mL of penicillin and streptomycin). The fibroblasts were removed using FibrOut^TM^, Fibroblast Growth Inhibitors, (#4-21501, CHI scientific) at 37°C, 5% CO2 incubator for 3 days. The EpCAM positive cell selection was performed by Magnetic-activated cell sorting (MACS) using mouse CD326 (EpCAM) MicroBeads (#130-105-958, Miltenyi Biotech). The KL-primary lung tumor EpCAM^+^ cells (JK-43P) and KL-metastatic lung tumor EpCAM^+^ cells (JK-43M) were cultured in complete culture medium and maintained at 37 °C and 5% CO2. EpCAM^+^, TTF1, KRAS^G12D^ and LKB1 expression level were confirmed by western blot. The lung and metastatic draining lymph nodes from JK-43 mouse were stained with H & E and scored by a board-certified lung pathologist using criteria previously published in ref. (Cell. 2014;159(2):440–455.)

### Allograft animal experiment

KL-GEMM derived allograft experiments were approved by the Institutional Animal Care and Use Committee (IACUC) of Emory University. Seven-week old (about 20g body weight) female FVB/NJ mice (#001800) were ordered from Jackson laboratory. JK-43M cells (2.5 x 10^6^ cells in 30% Growth factor reduced Matrigel in 100μl of PBS) were injected subcutaneously in the right flank of each mouse. Mice (n-5/group) received the following treatment: vehicle control and LDN214117 formulated in 2% DMSO + 30% PEG400 + 2% Tween 80 + ddH2O (25mg/kg, I.P. once daily) for 22days. Tumor size and body weight were measured twice a week.

### Immunohistochemistry

Formalin-fixed paraffin-embedded mouse tumor sections were stained with rabbit anti-BMP6, rabbit anti-hepcidin and rabbit anti-transferrin receptor antibodies. Secondary antibody kit (MP-7401-50) and HRP-based DAB Substrate kit (SK-4105) were purchased from Vector Laboratories and followed manufacturer’s instructions.

### Statistical analysis

A two-tailed unpaired Student’s t-test was used to analyze statistical significance between two conditions in an experiment. For experiments with three or more comparisons, an ordinary one-way ANOVA with a Tukey’s multiple comparisons test was used. Significance was assigned to p values <0.05; *p < 0.05, **p < 0.01, ***p < 0.001, ****p < 0.0001. Error bars represent the mean ± SEM.

## Acknowledgements

Research reported in this publication was supported by the National Cancer Institute of the National Institutes of Health under Award Number R01CA194027 (MGR, WZ, AIM), R01CA201340 (MGR, AIM), R01CA250422 (AIM), R01CA236369 (AIM), P50CA217691 (SSR, HF), P01CA257906 (HF, SSR), and Winship Cancer Institute Developmental Funds. Research reported in this publication was supported in part by the Emory Integrated Cellular Imaging Core, the Emory Integrated Genomics Core, the Emory Integrated Computational Core, the Cancer Animal Models Shared Resource of the Winship Cancer Institute of Emory University and NIH/NCI under award number P30CA138292 (SSR). The content is solely the responsibility of the authors and does not necessarily represent the official views of the National Institutes of Health.

---

**Figure S1.**
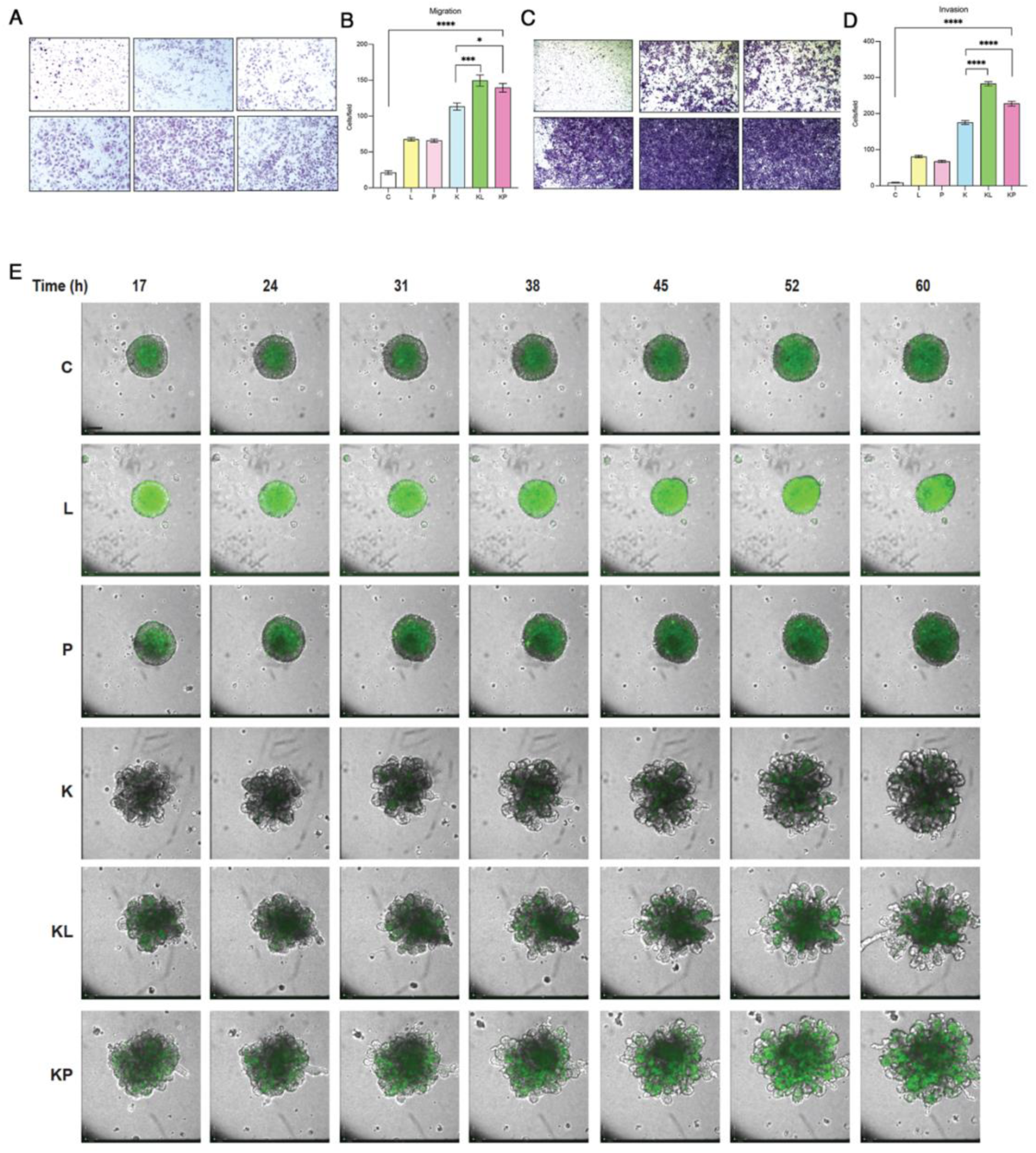
KL HBECs exhibit increased and phenotypically unqiue migration and early invasion patterns relative to other HBEC genotypes. A) Brightfield images of 2D migration assay from HBEC genotypes. B) Quantitation of 2D migration assay after 24hrs. C) Brightfield images of 3D invasion assay from HBEC genotypes. D) Quantitation of 3D invasion assay after 24hrs. Each genotype was run in triplicate and 10 fields/condition were counted. E) Brightfield and fluorescent images from 17hr-60 hour live-cell imaging of spheroids of the indicated genotype embedded in invasion matrix.

**Figure S2.**
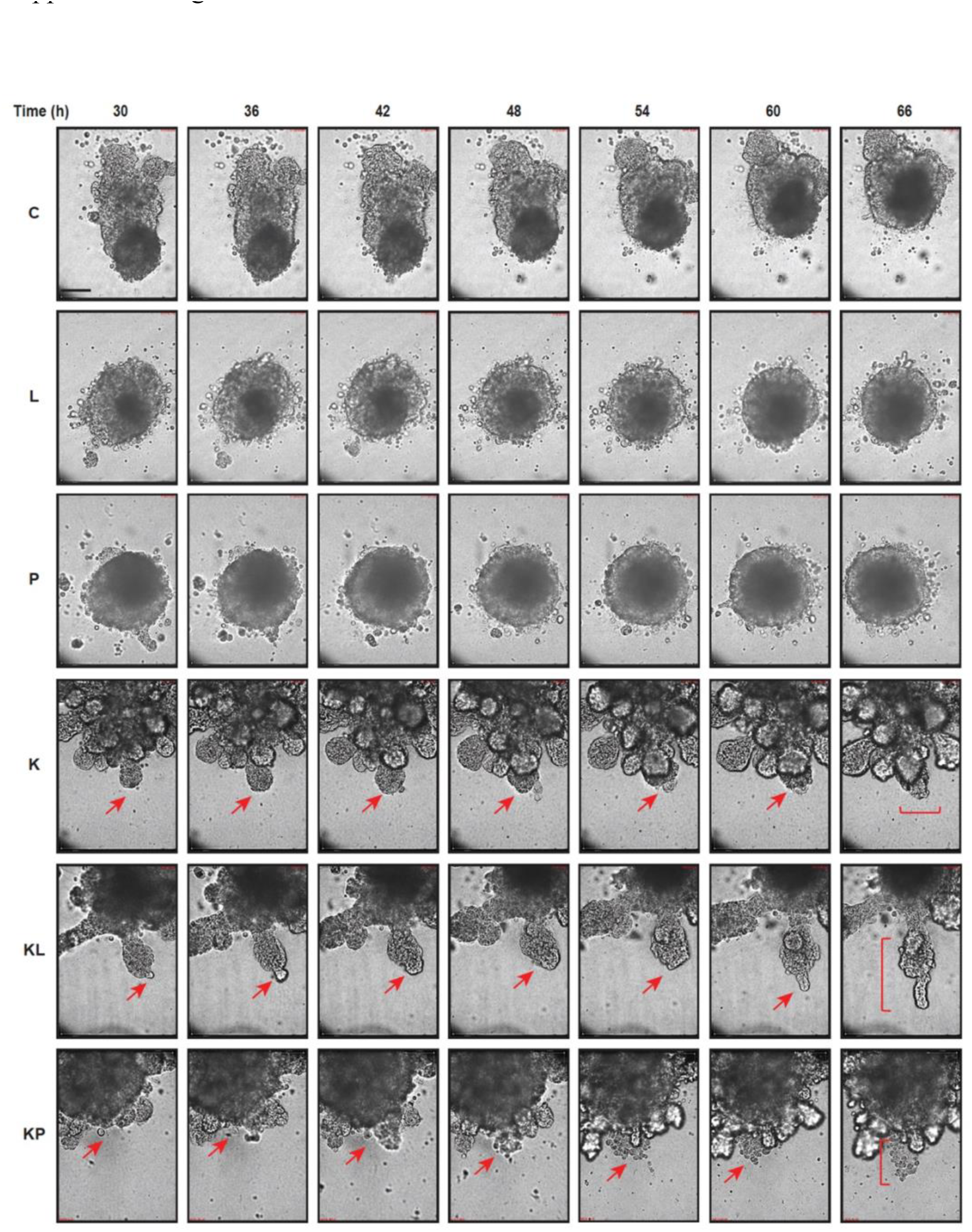
Isogenic HBECs exhibit subtype-specific behaviors in a 3D invasion matrix. Still images of HBEC spheroids of the indicated genotypes 30-66hrs post-embedding in invasion matrix. Red arrows and brackets indicate either branching morphogenesis (K), collective invasion (KL), or cell streaming (KP) phenotypes. Still images were extracted from Supplemental Movie 1.

**Supplemental Figure 3.**
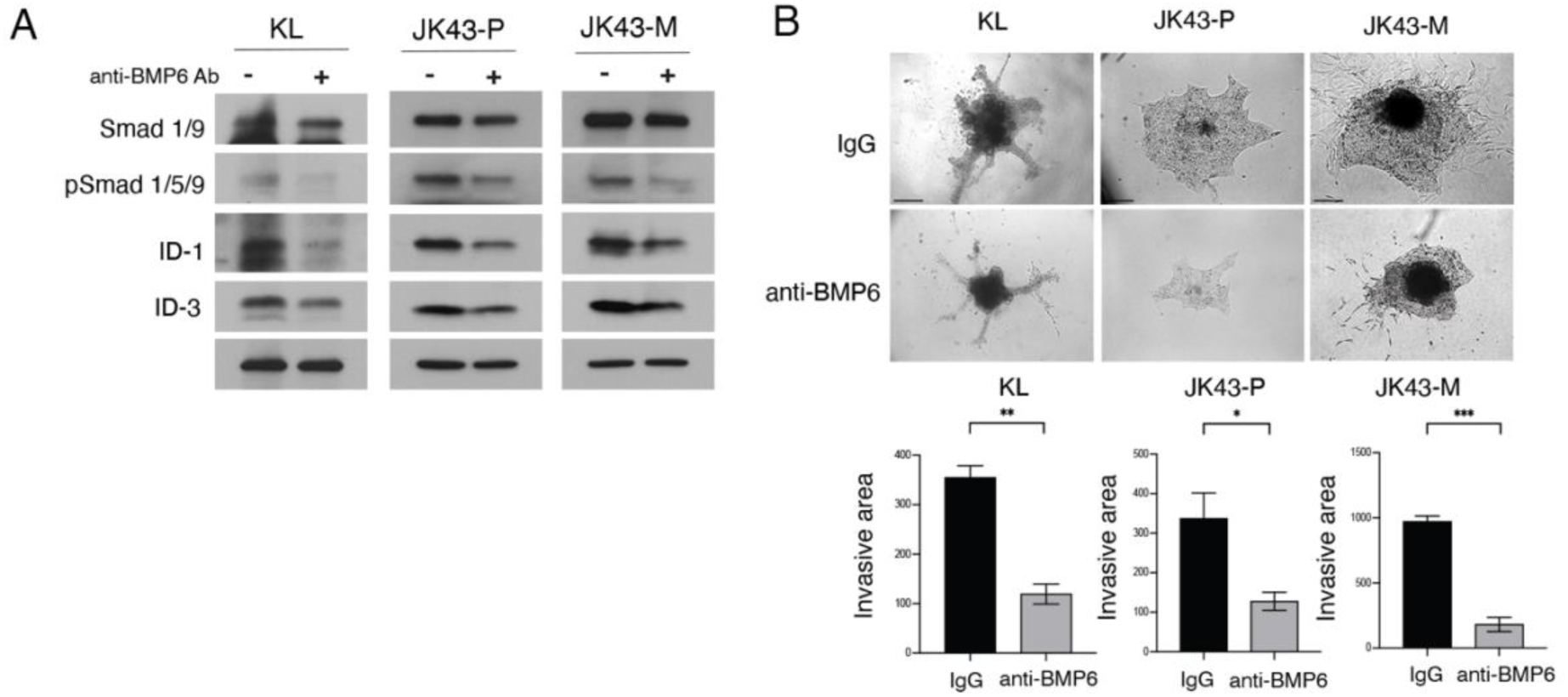
BMP6 neutralizing Ab attenuates Smad signaling and inhibits invasion in KL human and mouse cells. A) Western blot of cell lysates from KL HBECs, JK43-P and JK43-M cells treated with control or anti-BMP6 antibody and probed for the indicated signaling proteins. B) (top) Representative brightfield images of 3D invasion assay from the indicated cell lines treated with either IgG control or neutralizing anti-BMP6 antibody. Scale bar = 200 µm. Graph depicting mean invasive area for each experimental condition (bottom). Results are plotted as the mean of 3 independent experiments.

## References Cited

1. Salgia R, Pharaon R, Mambetsariev I, Nam A, Sattler M. The improbable targeted therapy: KRAS as an emerging target in non-small cell lung cancer (NSCLC). Cell Reports Medicine. 2021;2:100186.

2. Skoulidis F, Byers LA, Diao L, Papadimitrakopoulou VA, Tong P, Izzo J, et al. Co-occurring Genomic Alterations Define Major Subsets of KRAS-Mutant Lung Adenocarcinoma with Distinct Biology, Immune Profiles, and Therapeutic Vulnerabilities. Cancer discovery. 2015;5:860–77.

3. Xue JY, Zhao Y, Aronowitz J, Mai TT, Vides A, Qeriqi B, et al. Rapid non-uniform adaptation to conformation-specific KRAS(G12C) inhibition. Nature. 2020;577:1–5.

4. Palma G, Khurshid F, Lu K, Woodward B, Husain H. Selective KRAS G12C inhibitors in non-small cell lung cancer: chemistry, concurrent pathway alterations, and clinical outcomes. Npj Precis Oncol. 2021;5:98.

5. Carretero J, Shimamura T, Rikova K, Jackson AL, Wilkerson MD, Borgman CL, et al. Integrative Genomic and Proteomic Analyses Identify Targets for Lkb1-Deficient Metastatic Lung Tumors. CCELL. 2010;17:547–59.

6. Schabath MB, Welsh EA, Fulp WJ, Chen L, Teer JK, Thompson ZJ, et al. Differential association of STK11 and TP53 with KRAS mutation-associated gene expression, proliferation and immune surveillance in lung adenocarcinoma. Oncogene. 2016;35:3209–16.

7. Skoulidis F, Goldberg ME, Greenawalt DM, Hellmann MD, Awad MM, Gainor JF, et al. STK11/LKB1 Mutations and PD-1 Inhibitor Resistance in KRAS-Mutant Lung Adenocarcinoma. Cancer discovery. 2018;CD-18-0099.

8. Ndembe G, Intini I, Perin E, Marabese M, Caiola E, Mendogni P, et al. LKB1: Can We Target an Hidden Target? Focus on NSCLC. Frontiers Oncol. 2022;12:889826.

9. Miller AJ, Spence JR. In Vitro Models to Study Human Lung Development, Disease and Homeostasis. Physiology. 2017;32:246–60.

10. Kuang Y, Wang Q. Iron and Lung Cancer. Cancer Lett. 2019;464:56–61.

11. Sanchez-Duffhues G, Williams E, Goumans M-J, Heldin C-H, Dijke P ten. Bone morphogenetic protein receptors: Structure, function and targeting by selective small molecule kinase inhibitors. Bone. 2020;138:115472.

12. Vela D, Vela-Gaxha Z. Differential regulation of hepcidin in cancer and non-cancer tissues and its clinical implications. Exp Mol Medicine. 2018;50:e436.

13. Wohlhieter CA, Richards AL, Uddin F, Hulton CH, Quintanal-Villalonga À, Martin A, et al. Concurrent Mutations in STK11 and KEAP1 Promote Ferroptosis Protection and SCD1 Dependence in Lung Cancer. Cell Reports. 2020;33:108444–108444.

14. Mohedas AH, Wang Y, Sanvitale CE, Canning P, Choi S, Xing X, et al. Structure–Activity Relationship of 3,5-Diaryl-2-aminopyridine ALK2 Inhibitors Reveals Unaltered Binding Affinity for Fibrodysplasia Ossificans Progressiva Causing Mutants. J Med Chem. 2014;57:7900–15.

15. Raja E, Tzavlaki K, Vuilleumier R, Edlund K, Kahata K, Zieba A, et al. The protein kinase LKB1 negatively regulates bone morphogenetic protein receptor signaling. Oncotarget. 2016;7:1120–43.

16. Rooney L, Jones C. Recent Advances in ALK2 Inhibitors. Acs Omega. 2021;6:20729–34.

17. Carvalho D, Taylor KR, Olaciregui NG, Molinari V, Clarke M, Mackay A, et al. ALK2 inhibitors display beneficial effects in preclinical models of ACVR1 mutant diffuse intrinsic pontine glioma. Commun Biology. 2019;2:156.

18. Baranda J, Gordon MS, Parikh AR, Yang H, Pennock GK, Komarnitsky P, et al. Abstract CT134: Phase 1, first-in-human, dose-escalation, safety, pharmacokinetic (PK), and pharmacodynamic study of oral TP-0184, an activin receptor-like kinase-2 (ALK2) inhibitor, in patients (pts) with advanced solid tumors (ASTs). Cancer Res. 2022;82:CT134–CT134.

19. Williams E, Bagarova J, Kerr G, Xia D-D, Place ES, Dey D, et al. Saracatinib is an efficacious clinical candidate for fibrodysplasia ossificans progressiva. Jci Insight. 2021;6:e95042.

20. Konen J, Wilkinson S, Lee B, Fu H, Zhou W, Jiang Y, et al. LKB1 kinase-dependent and - independent defects disrupt polarity and adhesion signaling to drive collagen remodeling during invasion. Mol Biol Cell. 2016;27:1069–84.

21. Summerbell ER, Mouw JK, Bell JSK, Knippler CM, Pedro B, Arnst JL, et al. Epigenetically heterogeneous tumor cells direct collective invasion through filopodia-driven fibronectin micropatterning. Sci Adv. 2020;6:eaaz6197.

22. Gilbert-Ross M, Konen J, Koo J, Shupe J, Robinson BS, Wiles WG, et al. Targeting adhesion signaling in KRAS, LKB1 mutant lung adenocarcinoma. JCI insight. 2017;2:e90487.

